# Characterization of undocumented CO_2_ hydrothermal vent’s system in the Mediterranean Sea: implications for ocean acidification forecasting

**DOI:** 10.1101/2022.10.27.513997

**Authors:** Michela D’Alessandro, Maria Cristina Gambi, Cinzia Caruso, Marcella Di Bella, Valentina Esposito, Alessandro Gattuso, Salvatore Giacobbe, Martina Kralj, Francesco Italiano, Gianluca Lazzaro, Giuseppe Sabatino, Matteo Bazzarro, Lidia Urbini, Cinzia De Vittor

## Abstract

A previously undocumented shallow water hydrothermal field from Sicily (Southern Tyrrhenian Sea, Italy), is here described based on a multidisciplinary investigation. The field, covering an area of nearly 8000 m^2^ and ranging in depth from surface to −5 m, was explored in June 2021, to characterise the main physico-chemical features of the water column, describe bottom topography and features, and identify the main megabenthic and nektonic species. Twenty sites were investigated to characterize the carbonate system. Values of pH ranged between 7.84 and 8.04, ΩCa between 3.68 and 5.24 and ΩAr from 2.41 to 3.44. Geochemical analyses of hydrothermal fluids gases revealed a dominance of CO_2_ (98.1%) along with minor amounts of oxygen and reactive gases. Helium isotope ratios (R/Ra =2.51) and δ^13^C_CO2_ (3) support an inorganic origin of hydrothermal degassing of CO_2_ and the ascent of heat and deep-seated magmatic fluids to the surface. Visual census of fishes and megabenthos (mainly sessile organisms) allowed identification of 62 species, of which four are protected by the SPA/BIO Protocol and two by the International Union for Conservation of Nature. The macroalgae *Halopteris scoparia* and *Jania rubens* and the sponge *Sarcotragus* sp. were the dominant taxa in the area, while among fishes *Coris julis* and *Chromis chromis* were predominant. The preliminary description of this venting field indicates this site as an area of considerable interest and suitable for future experimental studies on ocean acidification.

## Introduction

Hydrothermal vents are worldwide recognized for their biological, ecological, cultural, and economic relevance [1, 2]. The upwelling of fluids makes them highly productive areas [3], which provide notably ecosystem services as provision, regulation and cultural services. Their relevant production is related to the high rates of chemosynthetic production by microbial communities that drew energy converting inorganic compounds such as H_2_S, CH_4_, and Fe^2+^ [4]. This high primary production leads to high biomass of invertebrates and fisheries resources, which in turn brings economic benefits to humans [3]. The benefits of hydrothermal vents also include the provision of metabolites and bioactive molecules produced by some organisms to survive in extreme conditions that are used by the cosmetic and pharmaceutical industries [5]. Moreover, the uptake of sulphide and methane by bacteria reduces the emission of these substances to the atmosphere [6]. In addition, only recently has been pointed out the role of vents in binding metals to organic molecules in seawater, creating organic-metallic complexes that increase their flux to the global ocean [2]. In terms of cultural services, the hydrothermal ecosystems represent interesting study areas in various scientific fields, including studies on the Ocean Acidification (OA) and the origin of life in the universe [7, 8]. In fact, in many vent’s systems, the bubbling of fluids makes the surrounding environment an ecological analogue of what is expected in future ocean scenarios as a consequence of OA [9, 10, 11]. For this reason, selected hydrothermal ecosystems are increasingly being used as natural laboratories to analyse species and community-level biological and physiological responses to OA [12, 13].

Hydrothermal fields are found in a wide depth range from the intertidal zone to the abyss and are characterized by different living communities [14, 15]. Generally, the deep hydrothermal vents (> 200 m) are inhabited by a typical fauna dominated by obligate taxa and symbiotic species which include siboglinid tubeworms, large bivalves, and snails [14]. The shallow-water (< 200 m) hydrothermal vent communities, which usually are not characterized by exclusive taxa are not substantially different from the neighbouring ones [15]. Despite their apparent normality, shallow vents have provided new or rare species, although not as closely related to hydrothermal fluids as in oceanic deep-waters vents [16, 17].

The biological peculiarities distinguishing deep and shallow hydrothermal areas are mainly tied to differences in their chemical complexity. In fact, shallow hydrothermal systems are characterized by lower pressure of fluids that may be derived from meteoric water, seawater, and magmatic fluid in variable amounts. The groundwater flow can be influenced by different sources, such as the hydraulic gradient, rock type, anthropogenic pressure, and tidal pumping [18]. In contrast, deep-sea hydrothermal fluids are derived exclusively from seawater, except in special cases where magmatic fluids are involved [19, 20]. To date, about 25 shallow systems are known worldwide [21, 22, 23], but a significant number may not yet have been recognized [24].

In the Mediterranean Sea, hydrothermal areas are reported from Tyrrhenian, Balearic and Aegean Sea. Tyrrhenian hydrothermal fields are known in the Aeolian archipelago, i.e., at Vulcano, Stromboli, Filicudi and Panarea Islands and nearby islets, at Capo Palinuro, Ischia island and other sites within the Bay of Naples (Campania), at the Pontine Archipelago (Latium), [22, 23, 25, 26], and Scoglio d’Africa (Tuscan Archipelago) [27]. In the Balearic Sea, a vent system has been described at the Columbretes islands [28], while in the Aegean, major hydrothermal vent systems are found along the Volcanic Arc at Euboea, Milos, Santorini, Kos, and Yali Nisiros [22, 29].

The recognition and characterization of hydrothermal vents are crucial for understanding the interactions between seafloor geochemical and biological processes. In particular, shallow hydrothermal vents ecosystems are of special interest due to their usually easy accessibility and suitability for experimental and manipulative research [14]. The aim of this paper is to provide the first multidisciplinary description of a previously undocumented CO_2_-rich hydrothermal system in the southern Tyrrhenian Sea (northern coast of Sicily), in order to characterize the area and discuss its suitability for experimental studies of ocean acidification.

## Material and Methods

### Geological setting of the investigated hydrothermal area

The studied hydrothermal area is located at San Giorgio di Gioiosa Marea, Gulf of Patti NE Tyrrhenian coast of Sicily (S1Table, Figs 1 and 2).

**Fig 1.**
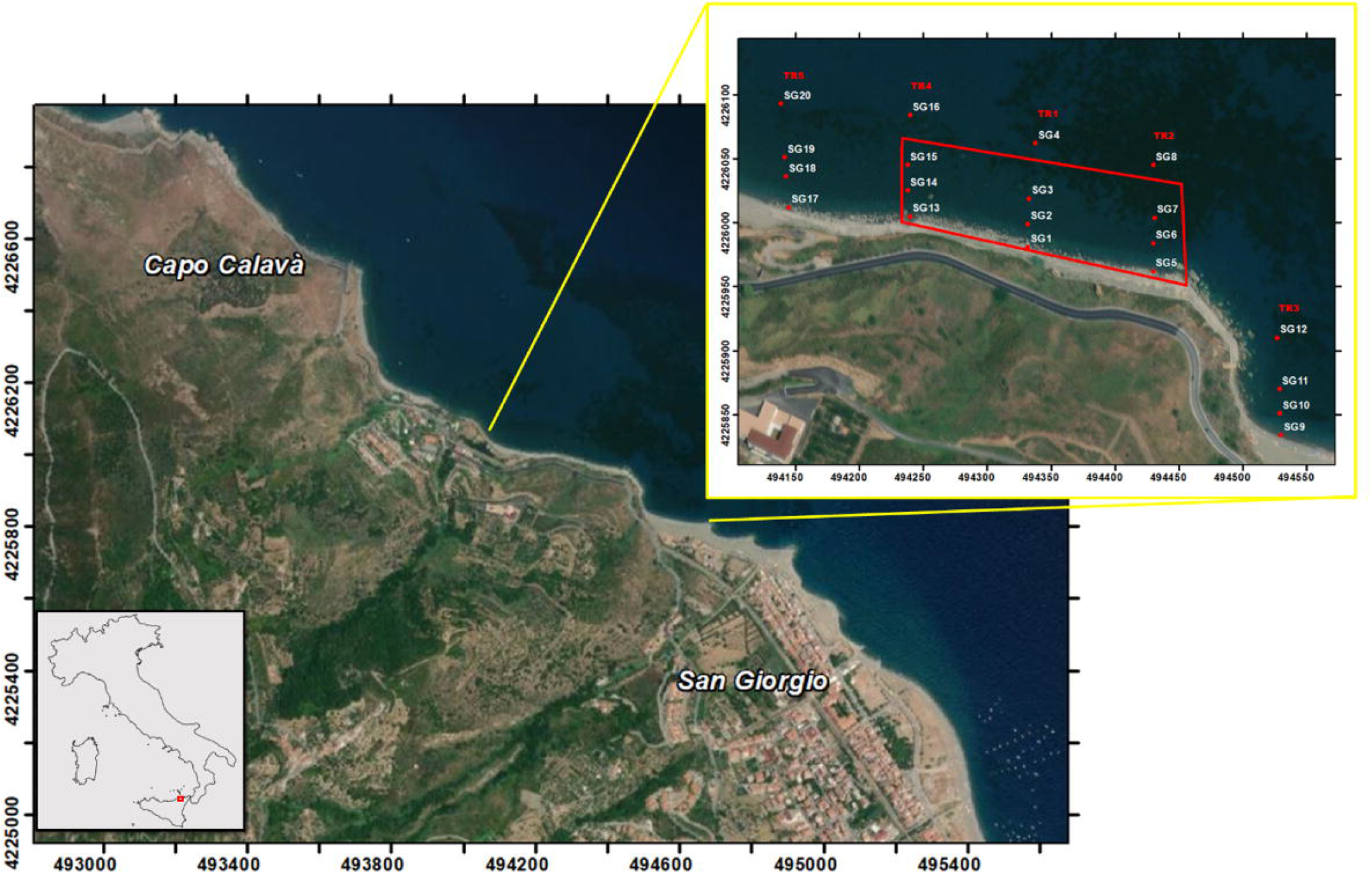
The study site and sampling spots. The hydrothermal field of San Giorgio is highlighted by the red polygon. Map made by means of ArcMap 10.5, ESRI software.

**Fig 2.**
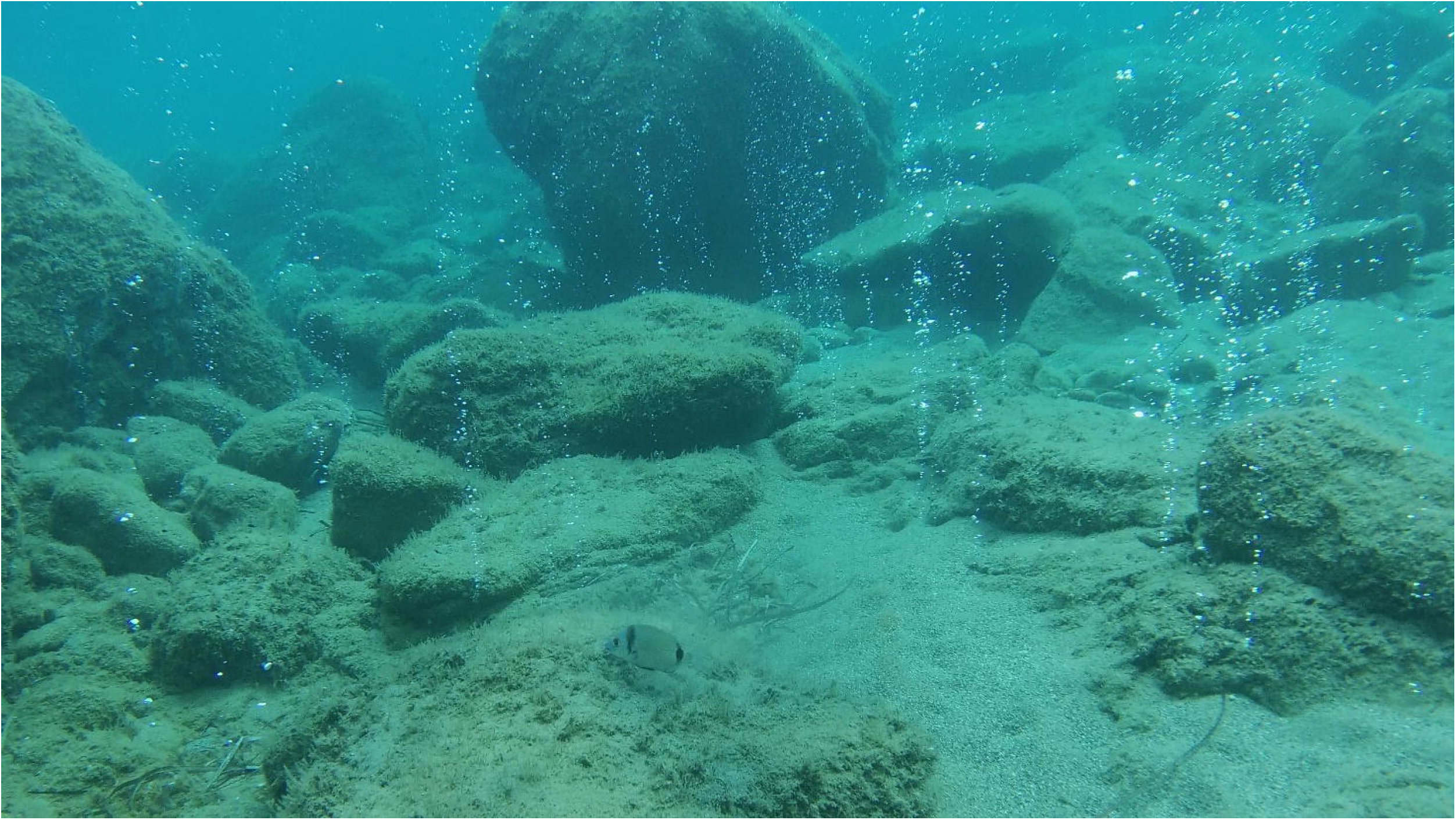
One of the high intensity gas emissions from the San Giorgio hydrothermal vents.

The area belongs to the regional tectonic structure known as the Aeolian-Tindari-Letojanni NW-SE faults system (ATLFS), which extends from the central sector of the Aeolian Islands (Salina, Lipari and Vulcano) to the Ionian Sea [30, 31, 32]. It represents one of the most seismically active regions of the Italian peninsula [31, 32] and is linked to the general geodynamic scenario of the central Mediterranean, characterised by the convergence between the European and African plates [33, 34, 35].

Recently, on the coast of the study area, CO_2_ degassing was detected by Italiano et al [26] along the southernmost section of the ATLFS (Nebrodi-Peloritani Mountains) suggesting a source in the crust and/or mantle for the fluid emissions in question. In addition, intense submarine volcanic processes and degassing activities have been reported in the nearby aeolian volcanic archipelago facing the study area, mainly on the islands of Vulcano and Panarea [36]. The CO_2_ vents of Panarea represent the major hydrothermal systems of the entire Mediterranean Sea, where a submarine gas eruption occurred in early November 2002 near the Bottaro rock, producing a submerged crater [37].

### Sampling survey

A preliminary survey was conducted in May 2021 to define the spatial distribution of the system, develop a sampling plan, and select the most appropriate sampling and investigation techniques. During this first step, a rectangular area approximately 200 m long and up to 40 m from the shoreline, covering the emission field, has been defined. Within this area, two different surveys were planned, respectively aimed to characterise the main physico-chemical features, and conduct a visual census to verify the bottom topography and features, and identify the main megabenthic and nektonic species.

Biological and chemical samplings were conducted in June 2021. Five transects perpendicular to the coastline were considered for seawater chemical analyses (Fig 1, S1 Table). A central transect (TR1) was established to intercept the point of highest emission flux at the bottom (SG2), moving eastwards, the TR2 crosses the last emissions visible and the TR3 transect is located about 100 m far from TR2. The other two transects were established to the west of TR1 near the last visible emissions (TR4) and about 100 m away from it (TR5), respectively. Four sampling stations (SG) were established in each transect at 1, 20, 40, and 80 m from the shoreline reaching 4 m of maximum depth, for a total of 20 stations, 9 of which within the hydrothermal field and 11 outside (transects TR3, TR5, and all sampling points 80 m from the shoreline). This sampling plan allowed us to evaluate the impact of emissions on the surrounding environment and the chemical differences among hydrothermal and not hydrothermal sites.

Biotic surveys were conducted using underwater visual census (UVC) in the area directly affected by the emissions (circumscribed by the red rectangle in Fig 1), using transects parallel to the shoreline to better assess local species diversity at similar depths. UVC observations were conducted in four transects along the shoreline length of the area (200 m) and spaced 10 m apart, covering an area of approximately 8000 m^2^ and a depth range of 1 (TR A) to 4-5 m (TR D), (S2 Table).

### Water chemistry analyses

Water samples for salinity, pH, total Alkalinity (A_T_), dissolved inorganic nutrients, and H_2_S determination were collected with Niskin bottles along the five transects perpendicular to the shoreline (TR1-TR5). Salinity was collected in 200-mL glass bottles, stored at 4 °C in the dark and then determined at the Centre for Oceanographic Calibration and Metrology (CTMO) of OGS by means of a Guildline Autosal 8400B Laboratory Salinometer.

For pH and AT samples, 120-mL borosilicate glass bottles were filled with seawater, poisoned with 100 μL of saturated mercuric chloride (HgCl_2_) to halt biological activity, sealed with glass stoppers and stored in the dark at 4 °C until analysis. Analyses of pH and AT were performed using a Cary 100 Scan UV-visible spectrophotometer and a Mettler Toledo G20 with LAUDA L100, respectively, according to the laboratory procedures described by Dickson et al [38] and Urbini et al. [39]. The analytical precision for pH was estimated to be ±0.002 pH_T_ units. Accuracy and precision of the AT measurements on CRM were determined to be less than ±2.0 mmol kg^-1^.

The samples for dissolved inorganic nutrients (nitrite, NO_2_, nitrate, NO_3_, ammonium, NH_4_, phosphate PO_4_, and silicic acid, H_4_SiO_4_) determination were filtered on 0.7 μm pore size glass-fibre filters (Whatman GF/F) and stored frozen (−20 °C) in polyethylene vials until laboratory analysis. Samples were defrosted and analysed colorimetrically using a segmented flow QUAATRO Seal Analytical AutoAnalyzer according to Hansen and Koroleff [40]. The detection limits for the method were 0.01 μM, 0.02 μM, 0.03 μM, 0.01 μM, and 0.01 μM for NO_2_, NO_3_, NH_4_, PO_4_, and H_4_SiO_4_, respectively. Samples for hydrogen sulphide determination (H_2_S) were collected in 40-mL vials adding 0.8 mL zinc acetate (ZnC_4_H_6_O_4_), stored at 4 °C in the dark and then measured spectrophotometrically using a VARIAN CARY 100 Scan spectrophotometer at 670 nm according to [41].

All the analyses were performed by the National Institute of Oceanography and Applied Geophysics (OGS) both at the laboratories in Trieste and at the ECCSEL-ERIC NatLab Italy in Panarea.

### Gas composition analyses

Gas analyses were performed to rule out the organic nature of the bubbling and to determine its hydrothermal features. Samples were collected from the submarine vents using an inverted funnel connected to two-way glass bottles [42]. The chemical composition of bubbling gases was determined by gas chromatography (GC) using an Agilent instrument with double TCD-FID detector and argon as carrier gas. Gas samples had been admitted to the GC by a syringe, and uncertainties are within ±5%. Measurements of the carbon isotopic composition (δ^13^C_CO2_) of the derived gasses were performed using the Delta Plus XP IRMS equipped with a Thermo TRACE GC and an interface to the Thermo GC/C III. The results (expressed in δ^13^C‰) are for the V-PDB (Vienna-Pee Dee Belemnite) standard, and the standard deviation of the ^13^C/^12^C ratio was ±0.2‰. The He isotope ratio (^3^He/^4^He) was analysed using a Helix SFT-Thermo static vacuum mass spectrometer after purification of He under high vacuum and cryogenic separation of Ne. The isotopic composition of helium is reported as R/ RA, namely, ^3^He/^4^He of the sample versus the atmospheric ^3^He/^4^He (RA = 1:386 × 10-6). Typical uncertainties are within ±5%.

To estimate the total gas output from the sediments, five-point measurements of the gas flux were made, calculating the emptying time of a 1-litre bottle connected to the bubbling point via an inverted funnel (Fig 3).

**Fig 3.**
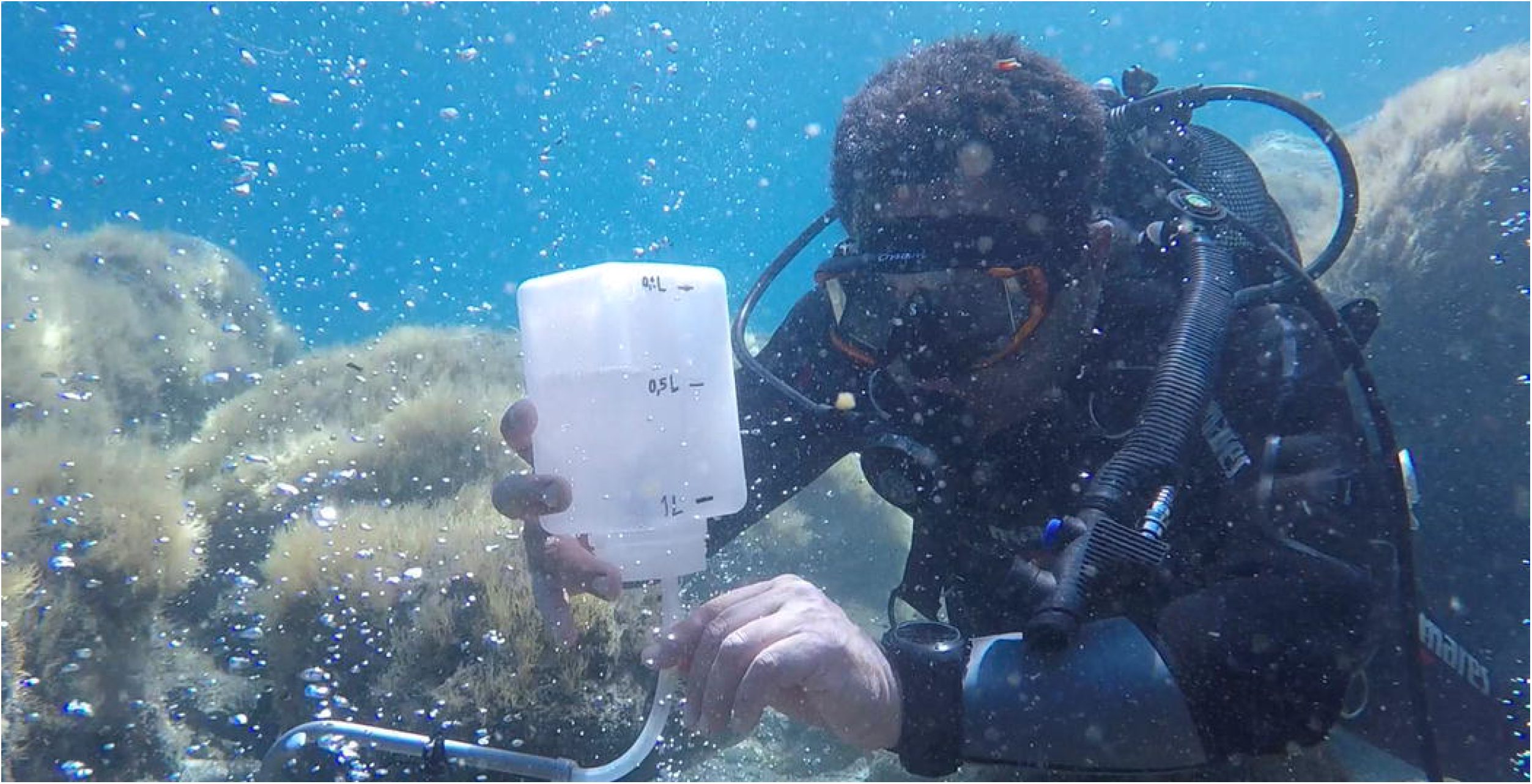
Measures of gas output flux from sediments by a INGV researcher at the San Giorgio vents.

These measurements were made in areas characterized by low, medium, and high fluxes, as defined by visual evaluation of the bubbling emission intensity by divers.

In addition, measurements of gas ejection concentrations were made using an innovative, ad hoc instrument for determining air-water gas exchange rates. It consists of a floating platform equipped with a specially designed infrared spectrophotometer (IR) directly connected to the chamber. The main components are an infrared gas analyzer (IRGA - CO_2_ Infrared Gas Sensor Gascard NG 10%), a pump and an accumulation chamber (S3 Figs). The chemical and isotopic analysis was performed in the laboratories of INGV-Palermo.

### Biotic investigation

Data on benthic and nektonic organisms and related bottom features were acquired through videos and photos from high-resolution cameras (Crosstour Action Cam 4K). The species directly identified in the field were recorded on a slate (S2 Table).

To calculate species and habitats frequency, videos of each transect parallel to the coast were analysed by selecting a frame every 10 seconds, corresponding to a linear distance of 2-3 m. In the selected plots (approx. 1×1 m frame), we estimated the percentage cover of different substrate types, such as soft sediments (sands), stones (lower than 1 m size), boulders (> 1 m size) and rocks, and the seagrass *Posidonia oceanica*; we also annotated in each plot the occurring benthic organisms (plant and animals) and fishes. The frequency of these organisms across the transect was measured as the percentage of their occurrence over the total plots examined (S2 Table). The scientific name of each identified species was up-dated according to the WoRMS database (https://www.marinespecies.org).

## Results

### Abiotic characterization

#### Water column chemistry

The results of the chemical characterization of the water column are shown in S1Table. With the exception of stations SG1 and SG12 (37.77 and 37.82, respectively), the salinity along all transects shows the constant value of 37.79.

Regarding to carbonate system parameters (Fig 4), A_T_ ranged from 2521 μmol kg^-1^, the minimum recorded at station SG3 (transect 1, depth 2.5 m), to 2533.5 μmol kg^-1^, the maximum measured at station SG4 (transect 1, depth 5 m). The pH_T_ at *in situ* temperature showed generally lower values along transects 1 and 4, reaching the minimum of 7.84 at station SG2. The other transects appeared to be less affected by vents and showed fairly homogeneous pH_T_ values, reaching the maximum (8.042) at station SG20. The lowest ΩCa out and ΩAr out were both recorded in SG2, with a value of 3.68 and 2.41, respectively.

**Fig 4.**
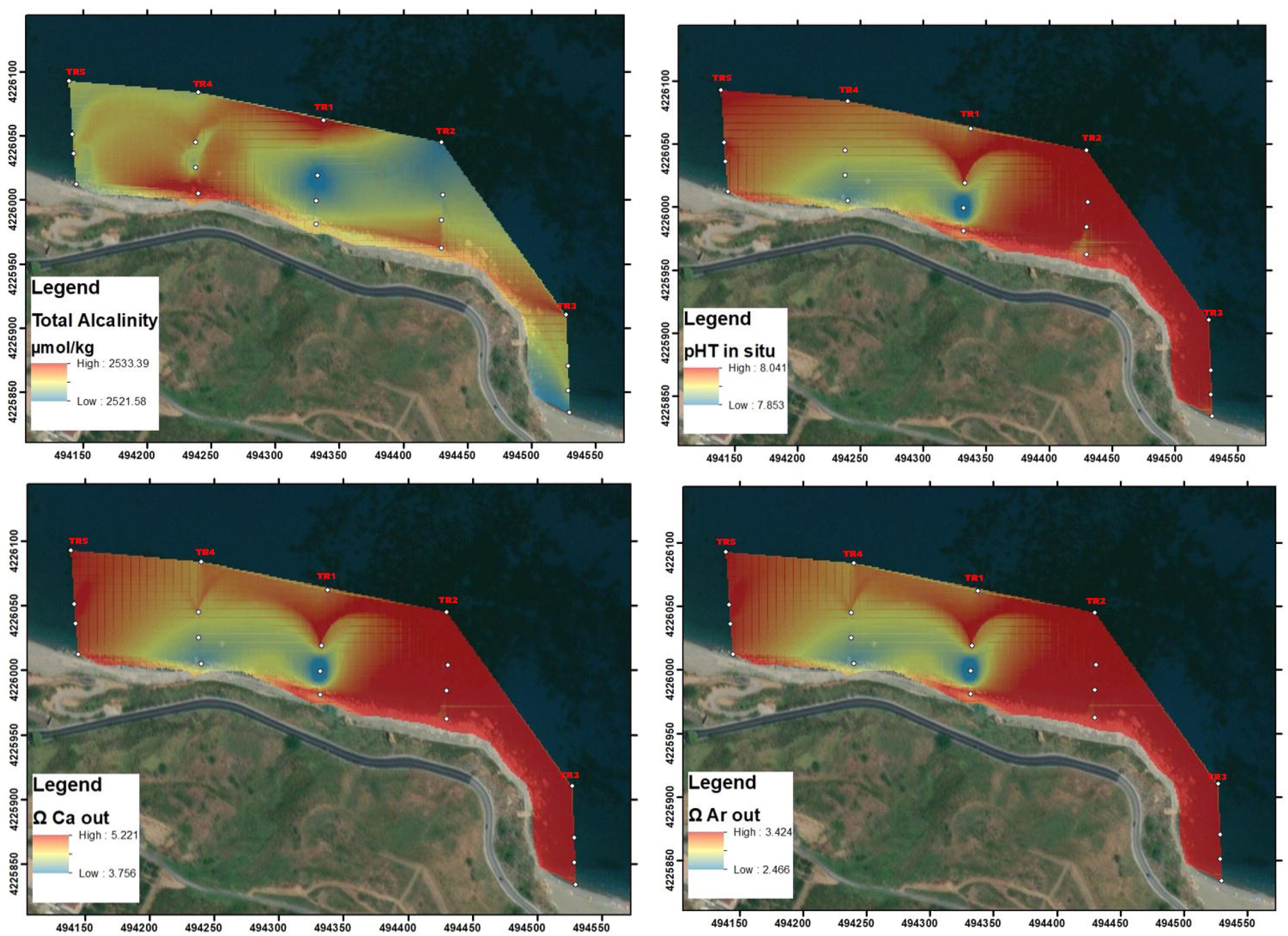
Natural Neighbour spatial interpolations of the main representative carbonate system recorded in the San Giorgio vents study area. Map made by means of ArcMap 10.5, ESRI software.

All dissolved inorganic nutrients, generally had the highest concentrations in TR5, outside of the direct bubbling of the vents on the westwards side (Fig 5 S1 Table). Ammonia ranged from undetectable (<0.03 μmol L^-1^, stations from SG10 to SG16) to 0.05 μmol L^-1^ (stations SG17 and SG18). Nitrite generally had concentrations of 0.01 – 0.02 μmol L^-1^ reaching the maximum of 0.04 in SG20. Nitrate showed very low concentrations along transects TR2, TR3 and TR4 (from undetectable to 0.04 μmol L^-1^), slightly elevated in TR1 (0.03-0.08 μmol L^-1^) and increased in TR5, reaching a maximum of 0.56 μmol L^-1^ at SG20.

**Fig 5.**
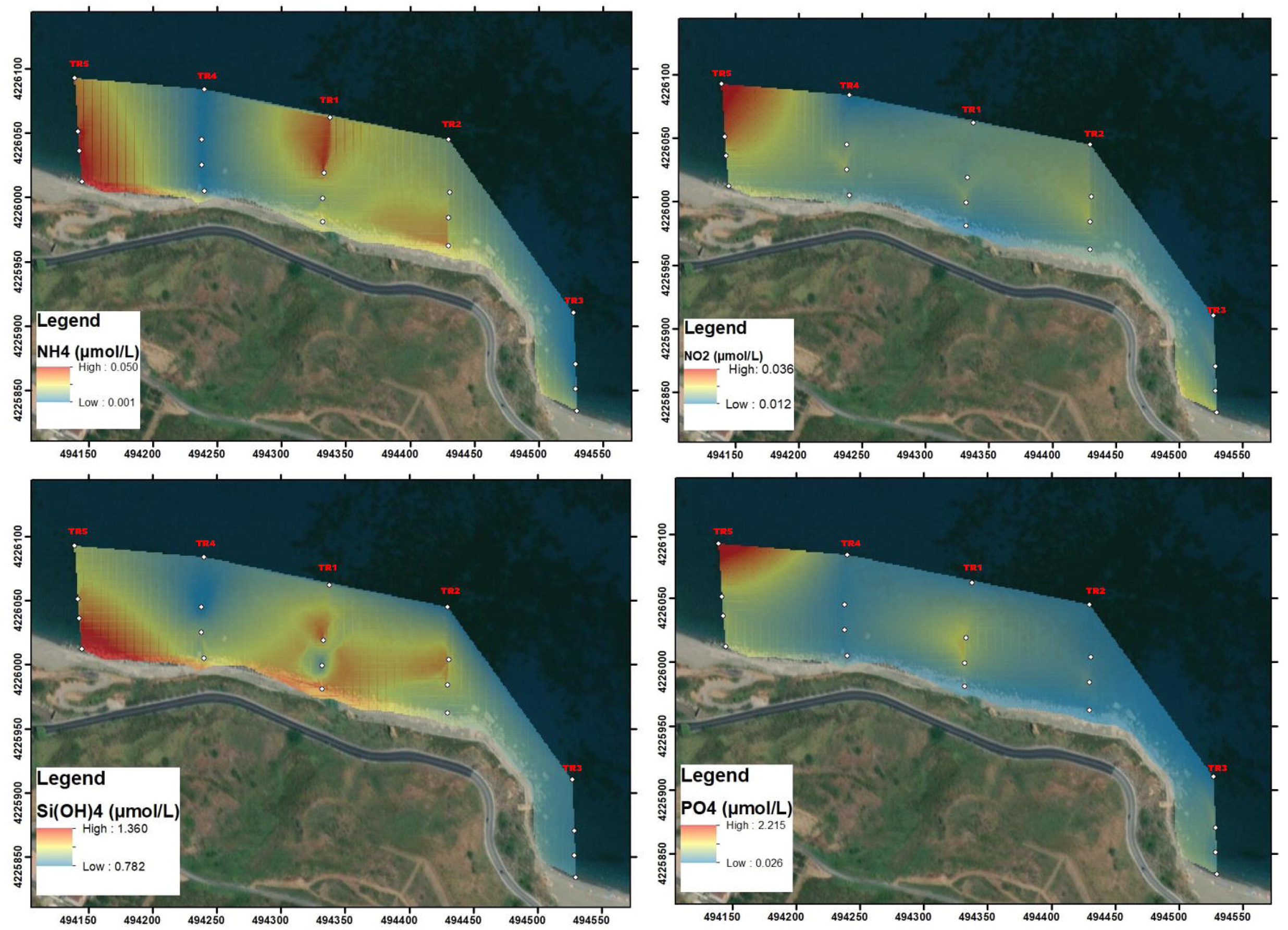
Natural Neighbour spatial interpolations of the main representative dissolved inorganic nutrients recorded in the San Giorgio vents study area. Map made by means of ArcMap 10.5, ESRI software.

Silicates ranged from 0.78 μmol L^-1^ in SG15 (TR4) to 1.36 μmol L^-1^ in SG17 (TR5), with a mean value of 0.96 ± 0.15 μmol L^-1^, while phosphates showed marked variations ranging from 0.02 μmol L^-1^ (SG5, TR2) to 2.24 μmol L^-1^ (SG17, TR5).

Spectrophotometric analysis of H_2_S showed the presence of this analyte, in low concentration (0.17 μM), only at station SG2, while it was undetectable at all other stations.

#### Gas characterization

Gas analysis revealed CO_2_ as the dominant gas component (98.1%), followed by N_2_ (1.02%), CH_4_ (0.43%), H_2_S (0.35%) and O_2_ (0.14%). In addition, very low percentages were found for He, H_2_ and CO (0.0015, 0.0007, and 0.0005%, respectively). The isotopic ratios of helium display values in the range of 2.51 Ra (Table 1).

**Table 1.** Chemical and isotopic compositions of the sampled gas emissions. concentrations are expressed in mol%. The helium isotopic composition is expressed as R/Ra, where R is the 3He/4He ratio in the sample and Ra is the same ratio in the atmosphere. The carbon isotopic composition is given versus PDB.

| R/Ra | He/Ne | $\delta^{13}\text{C}_{\text{CO}_2}$ | He % | $\text{H}_2$ % | $\text{O}_2$ % | $\text{N}_2$ % | CO % | $\text{CH}_4$ % | $\text{CO}_2$ % | $\text{H}_2\text{S}$ % |
| --- | --- | --- | --- | --- | --- | --- | --- | --- | --- | --- |
| 2.51 | 177.22 | 3 | 0.0015 | 0.000698 | 0.14 | 1.02 | 0.0005 | 0.43 | 98.1 | 0.35 |

The most abundant percentage of CO_2_ recorded at the air-water interface was of 1.73 % (Table 2).

**Table 2.** Percentage of CO_2_ recorded on four samples of air-water layer collected on the SG2 station. Coordinates are expressed in UTM WGS84, Zone 33 S.

| | North (m) | East (m) | $\text{CO}_2$ % |
| --- | --- | --- | --- |
| # 1 | 4226017 | 494330 | 0.7 |
| # 2 | 4226016 | 494331 | 1.17 |
| # 3 | 4226019 | 494332 | 0.85 |
| # 4 | 4226024 | 494335 | 0.35 |

A total flux of gasses of 30.75 litres/minutes has been estimated to be emitted in the whole area of the San Giorgio hydrothermal vents (Table 3).

**Table 3.** Total output of gasses and typologies of emission flux recorded on five different sites.

|  | Output<br>(litres/min) | Point<br>emissions | Measured<br>emissions | Total output<br>(litres/ min) | Typology<br>of emissions |
| --- | --- | --- | --- | --- | --- |
| **Flux 1** | 1 | 14 | 1 | 14 | High |
| **Flux 2** | 1.2 | 5 | 1 | 6 | High |
| **Flux 3** | 0.25 | 15 | 1 | 3.75 | Medium |
| **Flux 4** | 0.1 | 40 | 1 | 4 | Low |
| **Flux 5** | 0.1 | 30 | 1 | 3 | Low |

### Environmental description and biotic investigation

The seafloor, as observed along all transects parallel to the shoreline, consists of a mosaic of coarse sediments (mainly coarse sands) interspersed with stones (< 1 m), rocks/boulders (> 1 m), and patches of *Posidonia oceanica*. Sub-circular rusty patches occur on the sandy soil where intense venting occurs, ascribable to precipitation of metal oxides. The hard bottom areas (rocks or large boulders) close to the emissions are frequently colonized by microbial white mats. A total of 62 taxa were identified: 9 macrophytes, 29 invertebrates, and 24 fish (see list in S2Table).

Among the algae, all the species found were typical of the shallow photophilous hard bottoms, including *Padina pavonica, Halopteris scoparia, Anadyomene stellata, Ellisolandia elongata, Jania rubens* and *Codium bursa*. Among the invertebrates, the sponge *Sarcotragus* sp. is the most frequent sessile organism, while most of the other invertebrate species were molluscs (25%), mainly gastropods associated with vegetated hard bottoms, but also the bivalves *Chamelea gallina* and *Donax trunculus* were found, as associated with sandy substrate. Moreover, the bivalve *Pinna rudis* was recorded, associated with *Posidonia* meadows. Indeed, an extensive meadow of the seagrass *Posidonia oceanica* extends at the edge of the vent’s field. Two alien species were recorded, the green alga *Caulerpa cylindracea* and the crab *Percnon gibbesi*. The 24 species characterizing the ichthyofauna include *Coris julis*, *Chromis chromis*, *Diplodus annularis*, *Mullus barbatus*, *Thalassoma pavo*, *Trachinotus ovatus* and juveniles of two *Epinephelus* species, namely *E. costae* and *E. marginatus*, both considered as vulnerable and endangered in the IUNC list. Juveniles’ stages of several other bony fishes were recorded, such as *C. chromis*, *Diplodus vulgaris, M. barbatus*, *T. ovatus*, *T. pavo, Symphodus tinca* and *Sparisoma cretense*.

A brief description of each transect is provided below to characterize the main bottom features and the most frequent benthic and fish species.

Transect A (approx. 1 m depth, 109 plots analysed): the bottom is dominated by stones and sand, overall counting about the same frequency percentage of plots (82.6 and 81.7%, respectively).

Boulders/rock were recorded in 22% of plots. Very few tufts of *P. oceanica* were observed (2.7% of plots). Both boulders/rock and stone were characterized by a prevalent algal cover (92.7% of plots) mainly represented by turf and erected algae, with sparse sponges. The species recorded with the highest frequency was *Halopteris scoparia* (47.7% of the plots), followed by *Jania rubens* (28.4%), and by the sponge *Sarcotragus* sp. (14.7%). Several species were recorded with frequencies less than10%, as the coralline alga *Ellisolandia elongata* (7.3%), the sponges *Ircinia irregularis* (5.5%) and *Crambe crambe*, the green alga *Anadyomene stellata*, and the cnidarian *Pennaria disticha* (both with 2.8%). On the rocks and stones, the gastropods *Patella caerulea* (1.8%), *Stramonita haemastoma* (0.9%) and *Cerithium vulgatum* (0.9%) were also observed.

Gas emissions of different intensities occured in 5 scattered and 7 continuous plots. Along the transect, 12 bony fish species were observed, with *S. tinca* (17.4%) being the most frequent, followed by *C. chromis* (12%), and *D. vulgaris* (10%). The lowest frequencies were recorded for *E. costae* and *E. marginatus* (both with 0.9%).

In transect B (approx. 2 m depth, 96 plots analysed), the seafloor is composed of a mosaic of different substrate types: stones, occurring in 92.6% of the plots and covering a mean of 54% of the seafloor, followed by boulders/rock with 57% frequency and 50% covering, sands, although present in 82% of the examined plots, covered only a mean of 10% of the sea bottom. Stones showed a mean colonization by benthic organisms of 55%, while boulders/rock showed a mean colonization of 65%. *Posidonia oceanica* was present with very small patches in 22% of the plots and with a modest mean cover of 12%. Finally, gas emissions of different intensity were encountered in 13 scattered and 4 contiguous plots. Stones and rocks were mainly colonized by a turf of small filamentous algae (97% of plots), followed by 4 species of macroalgae such as *Padina pavonica* (78.1%), *J. rubens* (58.3%), *H. scoparia* (32.6%) and *A. stellata* (7.3%). Among invertebrates, only the sponge *Sarcotragus* sp. was dominant, occurring in 73% of the plots. Some gastropods were recorded with modest frequency, i.e., the sessile *Vermetus triquetrus* (3%), and the motile *Hexaplex trunculus* (4%). Seven taxa of bony fishes were observed, all with low frequencies: *C. chromis* (6.3%), *S. salpa* (4.2 %), *D. vulgaris* (3.1%), *O. melanura, S. cabrilla, T. pavo, Tripterygion* sp. (all with 2.1%).

On transect C (approx. 2.5-3 m depth; 85 plots examined) the seafloor still shows a mosaic of different substrates, but an increase in boulders/stones (64% of the plots) with higher mean bottom cover (59%), and a higher frequency of Posidonia patches was also observed, occurring in 62% of the plots and with a mean cover of 23.4%. Sand patches are still frequent (63%) although with modest mean cover (17%). Gas emissions were encountered in 3 scattered and 4 contiguous plots. At this depth, both stones and rocks show much higher levels of benthic colonization compared to transect B, with a mean of 66.8% for stones and 85% for boulders/rocks. The hard substrates continued to be colonized mainly by turf algae (95.5%) and by the previous cited four macroalgae, which increased their frequencies: *J. rubens* (72.9%), *P. pavonica* (89.4%), *H. scoparia* (71%), *A. stellata* (7.1%). Among the invertebrates still the most conspicuous species was *Sarcotragus* sp. (77.7%), followed by *V. triquetrus* (6%), the sponge *Crambe crambe* (3.4%), and colonial hydroids (likely *Eudendrium* sp.) (2.3%). Along the transect, 7 fish species were observed: *S. salpa* (5.9 %), followed by *D. vulgaris* (4.7%) and *O. melanura* (3.5%). Lower frequencies were recorded for *C. chromis, C. julis* (both 2.4%), *T. pavo* and *M. barbatus* (both 1.2%).

On transect D (approx. 4-4.5 m depth; 89 plots examined), the bottom is mainly characterized by sand (83.1%) mostly covered by *P. oceanica* (77.5%). Lower frequencies were recorded for stones (43.8%) and boulders/rock (7.9%), which showed high mean coverage percentages, 88.3% for boulders/rocks and 92.5% for stones. Only low intensity gas emissions were observed in 2 scattered and 4 continuous plots. The species showing the high frequencies were *P. pavonica* (33.7%), *J. rubens* (13.5%) *Sarcotragus* sp. (7.9%). Lower frequencies were recorder for the sponge *C. crambe* (1.1%) and the alga *A. stellata* (3.4 %). Six species of fish were encountered: *C. chromis* (21.3%), *S. maena* (7.9%), *O. melanura* (5.6%), *C. julis* and *D. vulgaris* (both with 2.2%), and *T. pavo* (1.1%).

## Discussions and conclusions

Shallow, CO_2_-rich hydrothermal vents provide unique opportunities to study the vulnerability of coastal ecosystems to ocean acidification [43]. In this context, the undocumented San Giorgio vents (southern Tyrrhenian Sea) is an appropriate case of study, suitable as a natural laboratory to study ocean acidification and related changes in benthic and fish population structure and habitat complexity, due to its physico-chemical features and easy accessibility [44]. Moreover, no other bubbling gasses and fluids were observed in the neighbouring areas during the study, although the presence of other emissions cannot be ruled out given the geological setting of the study area, which is located along the Aeolian-Tindari-Letojanni faults system [30, 31, 32]. In this area, the pH of 7.84 at the point of maximum emission flux, in the centre of the field indicated an intermediate level of acidification that is similar to the most reliable prediction for the year 2100 [45]. This value is comparable to those reported for the intermediate bubbling areas of the vents at the Castello Aragonese of Ischia [S2 and N_2_ stations of various publications, 14], whose pH, however, showed a great variability [46, 43] and a clear gradient, ranging from < 6.0 to 8.1, due to the high vents intensity in the most acidified zone, to normal pH conditions outside the vent area. At San Giorgio, the pH range is also similar to that reported for the lower pH zone in Baia di Levante at Vulcano (7.84 ± 0.24 pH) [48], as well as with the rim zone around Bottaro crater off Panarea island [49], in contrast with the very low values measured off Panarea around Basiluzzo island (pH 4.7-5.4) [50]. The contribution of the vents to the dissolved inorganic nutrient budget is apparently negligible, as suggested by the relatively low concentrations that increase in the outer part of the vent area, probably due to anthropogenic and terrigenous inputs.

The gasses emitted at the hydrothermal vents of San Giorgio have a predominant CO_2_ composition similar to those measured in Levante Bay (Vulcano Island) (97-99%) [48] and slightly higher than those measured in Castello Aragonese (90-95%) [14, 34]. The percentage of H_2_S (0.35%) in San Giorgio is lower than at Vulcano (2.2%) [51], while no sulphur occurs at the Castello Aragonese [14, 34]. This very low percentage of H_2_S, notably a toxic gas, favours investigations on the response of species and communities to ocean acidification. Indeed, in some vents the co-presence of CO_2_ with other toxic gasses or high metal ion concentration, makes it difficult to separate the negative effects of OA from those of other hazardous substances [52, 53]. The high R/Ra values measured in the San Giorgio vents could be related to the ascent of heat and deep magmatic fluids to the surface. These values are similar to those measured at Capo Calavà (R/Ra = 2.5) in a gas sample collected about 2 km WNW of the investigated area by Sano et al. [54]. The value of δ^13^C_CO2_ seems to confirm an inorganic origin of the hydrothermal degassing of CO_2_ [54]. Although no significant temperature differences were observed along the entire investigated area in our study, the observed sub-circular rusty patches, detected around the intense bubbling, could indicate an upwelling of mineral-enriched fluid columns into the seafloor, which needs to be further investigated.

Most species recorded in the San Giorgio vents are typical of shallow photophylous vegetated habitat and are commonly recorded in similar areas not subjected to gas emissions. The main algae dominating the rock and boulder coverage develop typically in summer (e.g., *P. pavonica, H. scoparia*), so that further important information would be acquired sampling at the vents in different seasons. However, the dominating taxon, along the whole transects and depths examined, is the perennial sponge *Sarcotragus* sp. Sponges are quite robust to OA, in accordance with their occurrence at other vent’s systems as at Castello Aragonese [55], Grotta del Mago cave [34], and Vullatura vents at Ischia; although in this latter system the community is dominated by *Crambe crambe* settled on dead matte of *Posidonia oceanica* [34]. Moreover, four species listed in Annexes I and II of the SPA/BIO Protocol of the Barcelona Convention were detected: the scleractinian *Cladocora caespitosa*, the sea urchin *Paracentrotus lividus*, the relative rare bivalve *Pinna rudis* and the seagrass *Posidonia oceanica*.

The fish fauna resembles that found in other analogous areas of the Mediterranean Sea, as reported for the vents at Castello on Ischia and Levante Bay on Vulcano [56]. Similarly to other shallow hydrothermal fields, no obligate taxa were identified in San Giorgio, whose communities were characterized by species, such as *S. salpa, C. chromis, C. julis*, that are known for their tolerance to acidification [56]. Although experimental studies have shown some negative effects of acidification on fish physiology and behaviour, especially on the juvenile stages [56, 57], the San Giorgio vent field would seem a nursery area, with occurrence of many juveniles of several species. In addition, juveniles of *Epinephelus costae* and *E. marginatus*, considered respectively as vulnerable and endangered species in the red list of threatened species enacted by the International Union for Conservation of Nature (IUCN), were also recorded.

Finally, differently than in other vents where alien organisms are often quite abundant, being favoured by low competition [25], a very low presence of alien species has been recorded in San Giorgio vent area, since limited to low covering by the alga *C. cylindracea* and sporadic occurrence of the decapod *P. gibbesi*,

In conclusion, the hydrothermal vent’s system of San Giorgio in Gioiosa Marea, characterized by predominant CO_2_ emissions and negligible presence of other potentially toxic compounds such as H_2_S, may be elected as a suitable natural laboratory to study the effects of OA on marine ecosystem processes, also benefiting from its easy accessibility and shallow depth.

Although hydrothermal vents are recognized as biodiversity hotspots and provide important ecosystem services, no protection measures have been proposed in the Mediterranean Sea. In the Natura 2000 network [58], habitat No. 1180, “Submarine structures made by leaking gases”, is considered as a habitat to be protected, but this definition is vague and seems to refer more to the geological nature rather than to the related biological aspects. We believe that the rarity of shallow hydrothermal vents and their ecological and evolutionary relevance require a better definition and explanation of 1180 habitat, bringing to include the biotic components associated with hydrothermal vents and their wide variability at local scales, in order to apply conservation measures and give more attention to these unique ecosystems.

## Acknowledgements

The authors wish to thank the Centre for Oceanographic Calibration and Metrology (CTMO) of OGS for providing salinity analyses and to Aqua Element Diving (San Giorgio, ME) for their technical assistance in scuba sampling activities, the IPANEMA HR project and the ECCSEL-ERIC - NatLab Italy managed by the OGS at Panarea, for logistic support.

## Supporting information

**S1 Table. Values of the physico-chemical variables measured in transects sampled perpendicular to the coastline off the San Giorgio vent area.**TR1= central transect intercept the points of highest bottom flow of emissions. TR2 = transect moving eastward in correspondence with the last emissions visible. TR3, about 100 m far from TR2. TR4= transect moving westward in correspondence with the last emissions visible. TR5= about 100 m far from TR4. In each transect four sampling stations (SG) were located at 1, 20, 40 and 80 m away from the shoreline, reaching 4.5 m of maximum depth.

**S2 Table. List of species visually recorded in the San Giorgio hydrothermal vent studied.**

Species without values of frequency for all transects are those recorded only during the explorative survey.

**S3 Fig 1.**
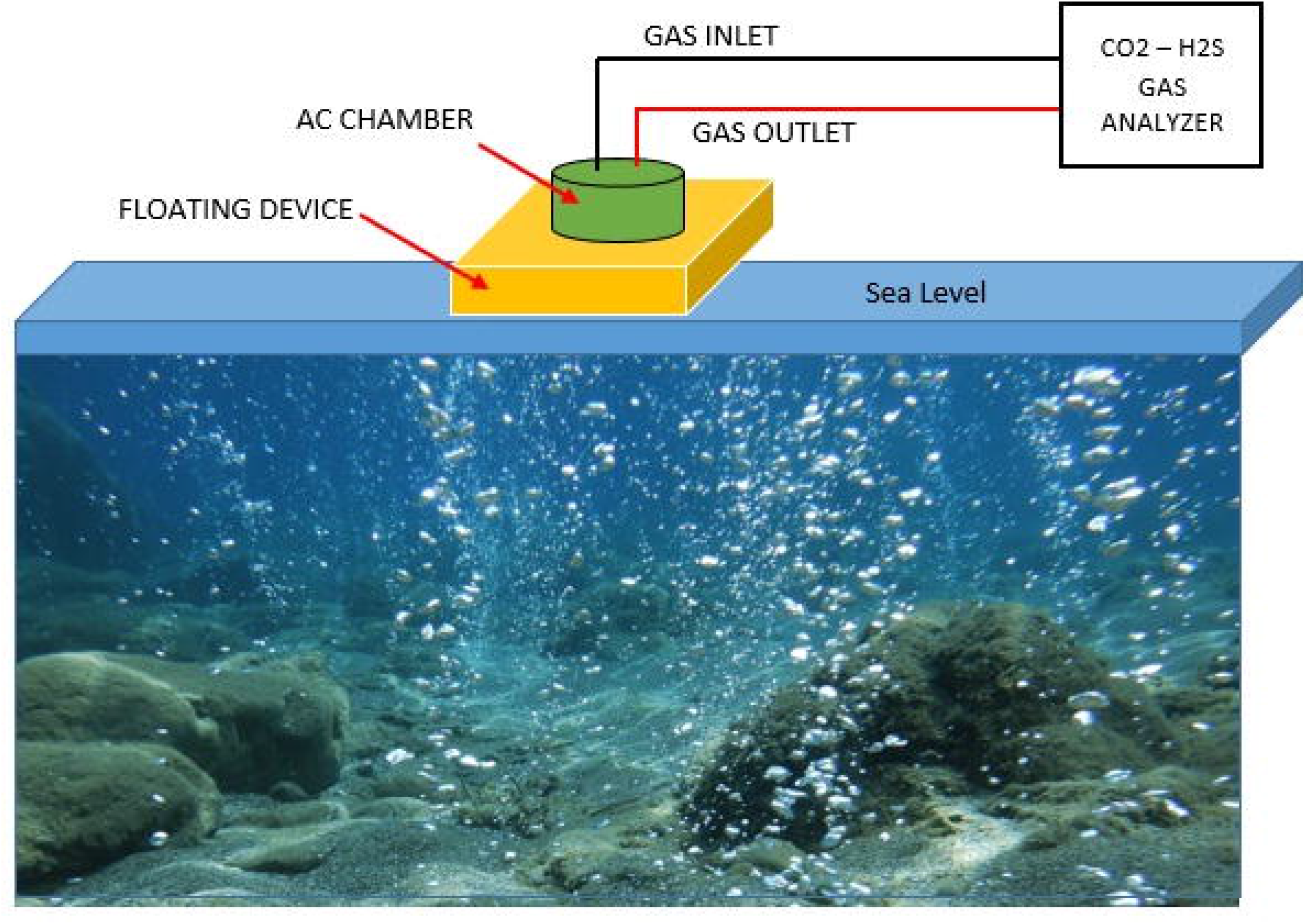
Draft of the Gas Output Instrument to measure the air-water gas exchange.

**S3 Fig 2.**
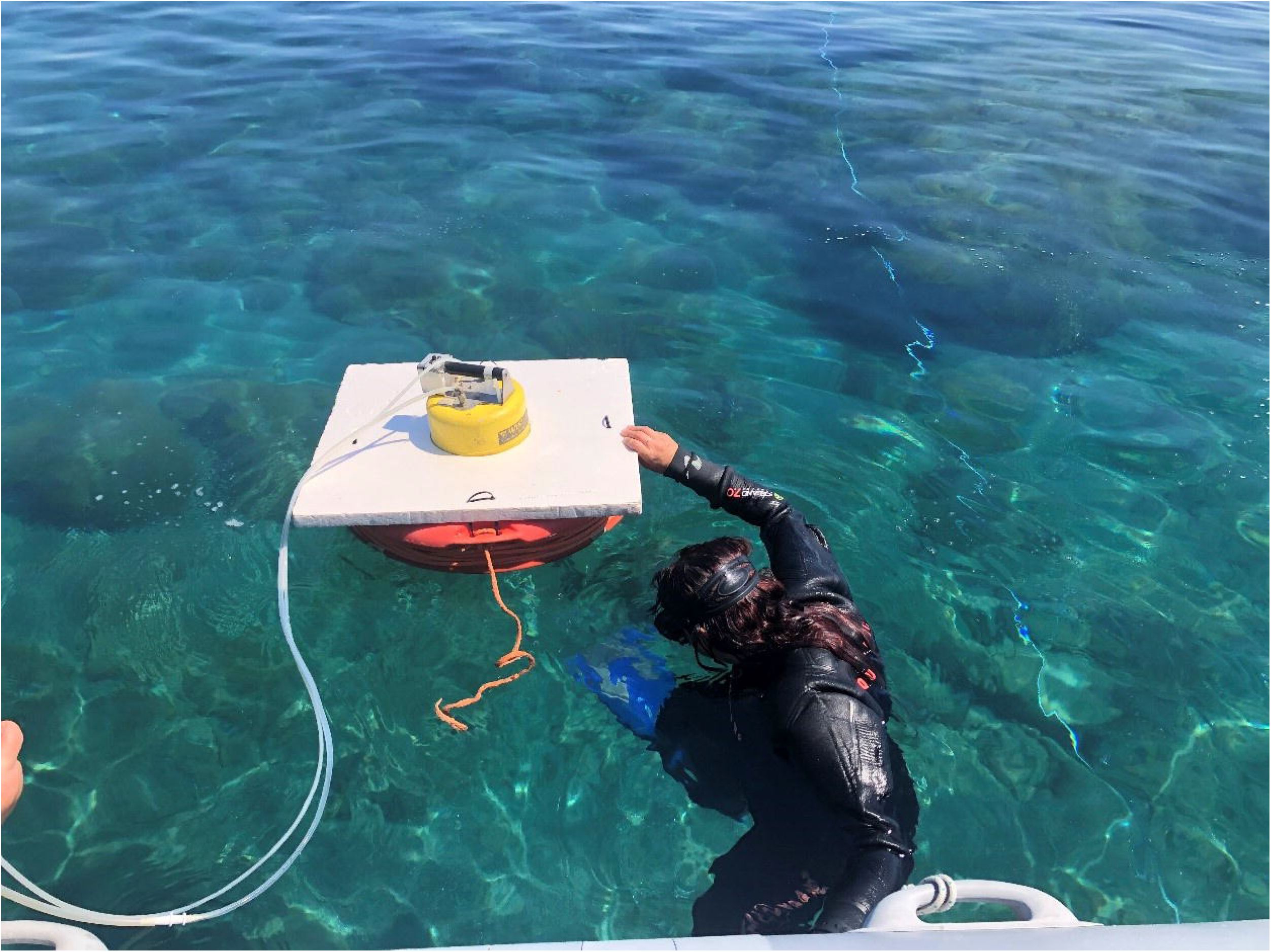
INGV researcher positioning the Gas output instrument on a sampling site off the San Giorgio vents.

