## Supplementary material for "Characterization of undocumented CO_2_ hydrothermal vent’s system in the Mediterranean Sea: implications for ocean acidification forecasting": S1

| **Transect** | **Station** | **Sampling date** | **Latitude N** | **Longitude E** | **Depth (m)** | **Distance from coast** | **Temperature in situ (°C)** | **Salinity** | **Total alkalinity (µmol/kg)** | **$pH_T$ *in situ*** | **$pCO_2$ out (µatm)** | **ΩCa out** | **ΩAr out** | **$NO_2$ (µmol/L)** | **$NO_3$ (µmol/L)** | **$NH_4$ (µmol/L)** | **$PO_4$ (µmol/L)** | **$Si(OH)_4$ (µmol/L)** |
|---|---|---|---|---|---|---|---|---|---|---|---|---|---|---|---|---|---|---|
| **TR1** | **SG1** | 4/6/2021 | 4225981 | 494332 | 0.5 | 0 | 22.1 | 37.77 | 2527 | 8.02 | 460 | 5.08 | 3.33 | 0.01 | 0.08 | <0.03 | 0.09 | 1.16 |
| **TR1** | **SG2** | 4/6/2021 | 4225999 | 494332 | 1.5 | 20 | 22.6 | 37.79 | 2524 | 7.84 | 748 | 3.68 | 2.41 | 0.02 | 0.07 | <0.03 | 0.70 | 0.84 |
| **TR1** | **SG3** | 4/6/2021 | 4226019 | 494333 | 2.5 | 40 | 22.2 | 37.79 | 2521 | 8.01 | 475 | 4.97 | 3.26 | 0.02 | 0.04 | 0.03 | 0.69 | 1.12 |
| **TR1** | **SG4** | 4/6/2021 | 4226062 | 494338 | 5 | 80 | 22.1 | 37.79 | 2534 | 7.98 | 516 | 4.73 | 3.10 | 0.02 | 0.03 | 0.03 | 0.11 | 0.91 |
| **TR2** | **SG5** | 4/6/2021 | 4225962 | 494430 | 0.5 | 0 | 22.6 | 37.79 | 2529 | 8.00 | 488 | 4.97 | 3.26 | 0.02 | 0.04 | 0.03 | 0.02 | 0.92 |
| **TR2** | **SG6** | 4/6/2021 | 4225984 | 494430 | 1.5 | 20 | 22.2 | 37.79 | 2527 | 8.01 | 473 | 5.01 | 3.28 | 0.02 | <0.02 | <0.03 | 0.28 | 1.04 |
| **TR2** | **SG7** | 4/6/2021 | 4226004 | 494431 | 2 | 40 | 22.1 | 37.79 | 2524 | 8.02 | 462 | 5.06 | 3.32 | 0.02 | 0.03 | <0.03 | 0.11 | 1.05 |
| **TR2** | **SG8** | 4/6/2021 | 4226045 | 494430 | 3 | 80 | 22.2 | 37.79 | 2523 | 8.03 | 450 | 5.16 | 3.38 | 0.02 | 0.02 | <0.03 | 0.08 | 0.81 |
| **TR3** | **SG9** | 4/6/2021 | 4225834 | 494530 | 0.5 | 0 | 22.6 | 37.79 | 2523 | 8.01 | 470 | 5.07 | 3.33 | 0.02 | 0.03 | <0.03 | 0.45 | 0.85 |
| **TR3** | **SG10** | 4/6/2021 | 4225851 | 494529 | 1.4 | 20 | 22.1 | 37.79 | 2527 | 8.04 | 438 | 5.24 | 3.44 | 0.01 | <0.02 | <0.03 | 0.12 | 0.84 |
| **TR3** | **SG11** | 4/6/2021 | 4225870 | 494529 | 2.7 | 40 | 22.2 | 37.79 | 2526 | 8.03 | 452 | 5.15 | 3.38 | 0.02 | 0.02 | <0.03 | 0.48 | 0.84 |
| **TR3** | **SG12** | 4/6/2021 | 4225910 | 494527 | 5 | 80 | 21.3 | 37.82 | 2529 | 8.03 | 452 | 5.02 | 3.29 | 0.01 | <0.02 | <0.03 | 0.03 | 0.85 |
| **TR4** | **SG13** | 4/6/2021 | 4226005 | 494240 | 0.5 | 0 | 22.4 | 37.79 | 2532 | 7.93 | 595 | 4.33 | 2.84 | 0.02 | <0.02 | <0.03 | 0.11 | 0.95 |
| **TR4** | **SG14** | 4/6/2021 | 4226025 | 494238 | 3.5 | 20 | 22.6 | 37.79 | 2526 | 7.95 | 556 | 4.54 | 2.98 | 0.02 | <0.02 | <0.03 | 0.13 | 0.93 |
| **TR4** | **SG15** | 4/6/2021 | 4226045 | 494238 | 2.9 | 40 | 22.3 | 37.79 | 2528 | 7.99 | 509 | 4.79 | 3.14 | 0.02 | <0.02 | <0.03 | 0.11 | 0.78 |
| **TR4** | **SG16** | 4/6/2021 | 4226084 | 494240 | 5 | 80 | 21 | 37.79 | 2525 | 8.01 | 478 | 4.79 | 3.13 | 0.01 | <0.02 | <0.03 | 0.15 | 0.84 |
| **TR5** | **SG17** | 4/6/2021 | 4226012 | 494145 | 0.5 | 0 | 22.5 | 37.79 | 2525 | 8.02 | 466 | 5.09 | 3.34 | 0.02 | 0.14 | 0.05 | 0.64 | 1.36 |
| **TR5** | **SG18** | 4/6/2021 | 4226036 | 494143 | 1 | 20 | 22.8 | 37.79 | 2528 | 8.02 | 469 | 5.13 | 3.37 | 0.02 | 0.09 | 0.05 | 0.60 | 1.16 |
| **TR5** | **SG19** | 4/6/2021 | 4226051 | 494142 | 2.8 | 40 | 22.7 | 37.79 | 2525 | 8.01 | 471 | 5.08 | 3.34 | 0.03 | 0.10 | 0.04 | 0.66 | 1.05 |
| **TR5** | **SG20** | 4/6/2021 | 4226093 | 494139 | 4.5 | 80 | 21.3 | 37.79 | 2527 | 8.04 | 435 | 5.14 | 3.36 | 0.04 | 0.56 | 0.03 | 2.24 | 0.98 |
