## Supplementary material for "Characterization of undocumented CO_2_ hydrothermal vent’s system in the Mediterranean Sea: implications for ocean acidification forecasting": S2

| | Percentage of frequency | | | |
|---|---|---|---|---|
| **ANIMALIA** | **TR A** | **TR B** | **TR C** | **TR D** |
| **Chordata** | | | | |
| *Aidablennius sphynx* (Valenciennes, 1836) | | | | |
| *Apogon imberbis* (Linnaeus, 1758) | | | | |
| *Chromis chromis* (Linnaeus, 1758) | 12 | 6.3 | 1.2 | 21.3 |
| *Coris julis* (Linnaeus, 1758) | 9.2 | | 2.4 | 2.2 |
| *Diplodus annularis* (Linnaeus, 1758) | | | | |
| *Diplodus puntazzo* (Walbaum, 1792) | | | | |
| *Diplodus sargus* (Linnaeus, 1758) | 4.6 | | | |
| *Diplodus vulgaris* (Geoffroy Saint-Hilaire, 1817) | 10 | 3.1 | 4.7 | 2.2 |
| *Epinephelus costae (*Steindachner, 1878) | 0.9 | | | |
| *Epinephelus marginatus* (Lowe, 1834) | 0.9 | | | |
| *Mullus barbatus* Linnaeus, 1758 | 5.5 | | 1.2 | |
| *Muraena helena* Linnaeus, 1758 | | | | |
| *Oblada melanura* (Linnaeus, 1758) | 2.8 | 2.1 | 3.5 | 5.6 |
| *Parablennius incognitus* (Bath, 1968) | 1.8 | | | |
| *Sarpa salpa* (Linnaeus, 1758) | | 4.2 | 5.9 | |
| *Serranus scriba* (Linnaeus, 1758) | 2.8 | | | |
| *Serranus cabrilla (Linnaeus, 1758)* | | 2.1 | | |

| | | | | |
|---|---|---|---|---|
| *Sparisoma cretense* (Linnaeus, 1758) | | | | |
| *Spicara maena* (Linnaeus, 1758) | | | | 7.9 |
| *Symphodus roissali* (Risso, 1810) | 4.6 | | | |
| *Symphodus tinca* (Linnaeus, 1758) | 17.4 | | | |
| *Thalassoma pavo* (Linnaeus, 1758) | 6.4 | 2.1 | 1.2 | 1.1 |
| Trachinidae gen. sp. | | | | |
| *Trachinotus ovatus* (Linnaeus, 1758) | | | | |
| *Tripterygion* sp. | | 2.1 | | |
| **Cnidaria** | | | | |
| *Balanophyllia (Balanophyllia) europaea* (Risso, 1826) | | | | |
| *Cladocora caespitosa* (Linnaeus, 1767) | | | | |
| *Pennaria disticha* Goldfuss, 1820 | 2.8 | | | |
| **Crustacea** | | | | |
| Balanidae gen. sp. | | | | |
| *Percnon gibbesi* (Milne Edwards, 1853) | | | | |
| **Echinodermata** | | | | |
| *Holothuria* sp. | 0.9 | | | |
| *Paracentrotus lividus* (Lamarck, 1816) | | | | |
| *Arbacia lixula* (Linnaeus, 1758) | | | | |
| **Mollusca** | | | | |
| *Cerithium vulgatum* Bruguière, 1792 | 0.9 | | | |
| *Chamelea gallina* (Linnaeus, 1758) | | | | |
| *Columbella rustica* (Linnaeus, 1758) | | | | |
| *Conus ventricosus* Gmelin, 1791 | | | | |
| *Donax trunculus* Linnaeus, 1758 | | | | |
| *Hexaplex trunculus* (Linnaeus, 1758) | | 4 | | |
| *Patella caerulea* Linnaeus, 1758 | 1.8 | | | |
| *Patella rustica* Linnaeus, 1758 | | | | |
| *Patella ulyssiponensis* Gmelin, 1791 | | | | |

| | | | | |
|---|---|---|---|---|
| *Phorcus articulatus* (Lamarck, 1822) | | | | |
| *Phorcus turbinatus* (Born, 1778) | | | | |
| *Pinna rudis* Linnaeus, 1758 | | | | |
| *Stramonita haemastoma* (Linnaeus, 1767) | 0.9 | | | |
| *Tarantinaea lignaria* (Linnaeus, 1758) | | | | |
| *Vermetus triquetrus* Bivona-Bernardi, 1832 | 3 | | 6 | |
| **Polychaeta** | | | | |
| *Serpula vermicularis* Linnaeus, 1767 | | | | |
| **Porifera** | | | | |
| *Crambe crambe* (Schmidt, 1862) | 2.8 | 1.0 | 3.4 | 1.1 |
| *Sarcotragus* sp. Schmidt, 1862 | 20.2 | 73 | 77.7 | 7.9 |
| *Ircinia irregularis* (Poléjaeff, 1884) | 5.5 | | | |
| **Ochrophyta** | | | | |
| *Padina pavonica* (Linnaeus) Thivy, 1960 | 14.7 | 78.1 | 89.4 | 33.7 |
| *Halopteris scoparia* (Linnaeus) Sauvageau, 1904 | 47.7 | 32.6 | 71 | |
| **PLANTAE** | | | | |
| **Rhodophyta** | | | | |
| *Ellisolandia elongata* (Ellis & Solander) Hind & Saunders, 2013 | 7.3 | 32.3 | 71.8 | |
| *Jania* cf *rubens* Lamouroux, 1812 | 28.4 | 58.3 | 72.9 | 13.5 |
| **Chlorophyta** | | | | |
| *Halimeda tuna* (Ellis & Solander) Lamouroux, 1816 | | | | |
| *Codium bursa* (Olivi) C. Agardh, 1817 | | | | |
| *Anadyomene stellata* (Wulfen) C. Agardh, 1823 | 2.8 | 7.3 | 7.1 | 3.4 |
| *Caulerpa cylindracea* (Sonder, 1845) | | 1.0 | | |
| **Tracheophyta** | | | | |
| *Posidonia oceanica* (Linnaeus) Delile, 1813 | 1.8 | | | |
