## Supplementary material for "Characterization of undocumented CO_2_ hydrothermal vent’s system in the Mediterranean Sea: implications for ocean acidification forecasting": S3

### Gas Output Instrument

The instrument is composed of three main parts (S3 Fig 1):

**S3 Fig 1. Draft of the Gas Output Instrument to measure the air-water gas exchange.**

1. Floating device: a platform allowing the chamber to float on seawater
2. AC chamber: accumulation chamber equipped with a pump that transfers gases to gas detector.
3. Gas analyser: $CO_2$ (IRGA - CO2 Infrared Gas Sensor Gascard NG 10%) and electrochemical sensors (0 - 100 ppm) for $H_2S$. During the measures, the chamber was sealed at the water surface, and continuous measurements of CO2 and $H_2S$ were made for approximately 3 min immediately after deployment (S3 Fig 2).

**S3 Fig 2. INGV researcher positioning the Gas output instrument on a sampling site off the San Giorgio vents.**

After a given time of pump activation, the value of gases concentration inside the probe reaches a steady value.
